# Pattern recognition receptors confer plant salt tolerance via WRKY18/WRKY40 transcription factors

**DOI:** 10.1101/2020.05.07.082172

**Authors:** Eliza P. Loo, Yuri Tajima, Kohji Yamada, Taishi Hirase, Hirotaka Ariga, Tadashi Fujiwara, Keisuke Tanaka, Teruaki Taji, Imre E. Somssich, Jane E. Parker, Yusuke Saijo

**Author notes:** Institute for Molecular Physiology, Heinrich-Heine-Universität, 40225 Germany (Eliza P. Loo). Graduate School of Technology, Industrial and Social Sciences, Tokushima University, Tokushima, 770-8501 Japan (Kohji Yamada). Leaf Tobacco Research Center, JAPAN TOBACCO Inc, Oyama, 323-0808 Japan (Taishi Hirase). Genetic Resources Center, NARO, Ibaraki, 305-8602 Japan.

## Abstract

Pattern recognition receptors (PRRs) bind microbe- and damage-associated molecular patterns (MAMPs/DAMPs, respectively) to enhance host immunity in animals and plants. Here, we report that PRRs also confer salt tolerance in the model plant *Arabidopsis thaliana* following recognition of cognate ligands, such as bacterial flagellin and EF-Tu and the endogenous Pep peptides. Pattern-triggered salt tolerance (PTST) requires the PRR-associated kinases *BAK1* and *BIK1*, and the NADPH oxidase *RBOHD*. Transcriptome profiling reveals an inventory of PTST target genes, which increase or acquire salt responsiveness following an exposure to immunogenic patterns. In their regulatory DNA sequences, specific binding sites for a subset of WRKY transcription factors are over-represented. Accordingly, PTST requires *WRKY40* and *WRKY18*, which activate salt tolerance-related genes but attenuate pathogen defense-related genes, including the *EDS1* immunity activator. PRR signaling leads to sustained WRKY40/WRKY18 accumulation under salt stress and utilizes both *WRKY*s for salt tolerance. The PRR-WRKY40/WRKY18 module also confers salt tolerance after challenge with non-pathogenic bacteria. Our findings give molecular insight into signaling plasticity underlying biotic-abiotic stress cross-tolerance in plants conferred by PRRs.

## INTRODUCTION

Like animals, plants have evolved an elaborate immune system to sense and adapt to disturbance caused by biotic agents. How the immune system influences abiotic stress responses remains much less understood. Plants sense and cope with fluctuating environments, while accommodating a rich diversity of microbial communities that often aid host adaptation. Conversely, environmental abiotic factors, such as light, temperature and water availability profoundly influence the mode and outcome of plant-microbe interactions (Velásquez *et al*, 2018). This predicts an intimate relationship between biotic and abiotic stress sensing and signaling in plants. In line with this, it is becoming apparent that immune receptors and signaling regulators also impact abiotic stress responses, positively or negatively in a context-dependent manner (Saijo & Loo, 2020). However, the regulatory logic or molecular basis behind intricate cross-regulations between biotic and abiotic stress signaling remains poorly understood.

Plant immunity largely relies on two classes of innate immune receptors, namely cell surface-localized PRRs and intracellular nucleotide-binding domain and leucine-rich repeat (LRR)-containing receptors (NLRs) (Jones & Dangl, 2006). Detection of MAMPs and DAMPs by cognate PRRs leads to pattern-triggered immunity (PTI), which is vital in preventing the infection of most non-adapted microbes and in restricting growth of adapted microbes, termed basal resistance (Boller & Felix, 2009; Couto & Zipfel, 2016). In turn, plant-infecting microbes, whether pathogenic or non-pathogenic, employ an array of effectors to manipulate host immunity and other processes for infection. To counter this, plants employ a repertoire of NLRs that recognize microbial effectors to mount effector-triggered immunity (ETI) that terminates microbial growth. NLRs are classified into two major subclasses, based on their N-terminal domains: the coiled-coil (CC)-NLRs and the Toll-interleukin1-receptor (TIR)-NLRs. CC-NLR and TIR-NLR functions typically require the defense regulators NDR1 and EDS1, respectively (Jones *et al*, 2016). Functional linkage between PTI and ETI through the actions of microbial effectors is illustrated by the so-called “zigzag model”, a principle that broadly applies across a wide range of plant-microbe interactions (Jones & Dangl, 2006). Compared to PTI, ETI is typically greater in amplitude and robustness against microbial perturbations, and is often accompanied by localized cell death called the hypersensitive response (HR) (Cui *et al*, 2015).

A major class of PRRs are the LRR-receptor kinases (RKs), including FLS2, EFR and PEPR1/PEPR2, which recognize bacterial flagellin (flg22 epitope), elongation factor Tu (EF-Tu, elf18 epitope) and the endogenous Pep epitopes embedded in their precursors, PROPEPs, respectively (Gómez – Gómez & Boller, 2000; Zipfel *et al*, 2006; Yamaguchi *et al*, 2006, 2010; Krol *et al*, 2010). Following ligand binding, these PRRs form heteromeric receptor complexes with the LRR-RK BAK1 (and related SERKs), and then induce dissociation of receptor-like cytoplasmic kinases (RLCKs) such as BIK1 and PBL1. Their trans-phosphorylation provides a basis for intracellular defense signaling, which involves Ca^2+^ release and an RBOHD-dependent reactive oxygen species (ROS) burst, phosphorylation cascades of Ca^2+^-dependent protein kinases and mitogen-activated protein kinases (MAPKs), callose deposition, production of the phytohormones ethylene and salicylic acid (SA), and extensive reprogramming of the transcriptome and proteome (Couto & Zipfel, 2016; Yu *et al*, 2017; Saijo *et al*, 2018). These signaling events collectively contribute to PTI, and also provide possible internodes for balancing immunity and other cellular processes.

PTI activation involves the transcriptional reprogramming of thousands of genes in *A. thaliana*, including transcription factor gene families (Birkenbihl *et al*, 2017b). The WRKY transcription factor family harbor a WRKY DNA-binding domain to bind W-box-containing cis-acting regulatory DNA elements in the target genes, thereby positively or negatively regulating their transcription (Rushton *et al*, 2010). Over-representation of the W box in the regulatory DNA sequences of flg22-inducible genes points to the extensive engagement of the WRKY family in PTI (Navarro, 2004). Indeed, many *WRKY* members are induced during PTI, including structurally-related *WRKY18* and *WRKY40* that negatively regulate PTI and resistance against biotrophic fungal pathogens and insects (Xu *et al*, 2006; Pandey *et al*, 2010; Lozano-Durán *et al*, 2013; Schön *et al*, 2013; Schweizer *et al*, 2013). Genome-wide identification of DNA binding sites and transcriptionally controlled targets indicates that WRKY18/WRKY40 attenuate early defense-inducible genes during PTI (Birkenbihl *et al*, 2017a). Conversely, *WRKY18/WRKY40* contribute positively to ETI via the TIR-NLR *RPS4* (Schön *et al*, 2013), and to necrotrophic fungal resistance (Xu *et al*, 2006). *WRKY18/WRKY40* have also been implicated in abiotic stress responses, since mutation of *WRKY18* enhances salt/osmotic tolerance in a manner requiring *WRKY40* (Chen *et al*, 2010; Shang *et al*, 2010). WRKY18, WRKY40 and related WRKY60 constitute the Group IIa subclade of the WRKY family (Rushton *et al*, 2010). These three *WRKY* members negatively influence ABA-mediated inhibition of seed germination and seedling growth (Yan *et al*, 2013). Their possible contributions to combined stress responses are unclear.

Amplification of PRR-triggered signaling is closely associated with effective pathogen resistance (Lu *et al*, 2009; Tsuda *et al*, 2009; Serrano *et al*, 2012). SA is a key for this process in biotrophic/hemibiotrophic pathogen resistance, and is produced in large part through the SA biosynthetic enzyme isochorismate synthase1 (ICS1) during PTI (Wildermuth *et al*, 2001; Vlot *et al*, 2009). SA signaling relies on the SA-binding transcriptional co-activator NPR1 and co-repressors NPR3/NPR4 (Ding & Ding, 2020), and also on EDS1 and related PAD4 (Wiermer *et al*, 2005). EDS1/PAD4 activate *ICS1* expression and SA accumulation but also promote *ICS1*/SA-independent defenses (Glazebeook *et al*, 2003; Bartsch *et al*, 2006; Cui *et al*, 2017). Accordingly, *EDS1* is required for basal resistance to biotrophic and hemi-biotrophic pathogens (Wiermer *et al*, 2005). However, excessive de-repression of *EDS1/PAD4-mediated* defenses during osmotic stress results in a collapse of osmotic stress tolerance (Ariga *et al*, 2017). Therefore, tight control of EDS1/PAD4 activity is crucial not only under biotic but also abiotic stress conditions.

Genetic studies have implicated PRRs in salt stress tolerance. In *Arabidopsis thaliana*, ectopic expression of fungal chitinase or chitin application enhances salt tolerance in a manner dependent on the lysin-motif (LysM) RK *CERK1*, which mediates the perception of fungal chitin and bacterial peptidoglycans (Brotman *et al*, 2012). Even under sterile conditions in the absence of microbes or MAMPs, *cerk1* plants are hypersensitive to salt stress (Espinoza *et al*, 2017). These studies suggest that CERK1 has a role in promoting salt stress tolerance, and that this function is related to an as-yet-unidentified endogenous DAMP(s). Likewise, *PROPEP3* overexpression and Pep3 application under sterile conditions both enhance salt tolerance through *PEPR1* (Nakaminami *et al*, 2018). These studies suggest that DAMP sensing and signaling contribute to salt stress tolerance, yet the underling mechanisms are not defined.

Here, we report that PTI signaling components promote salt tolerance in *Arabidopsis thaliana* following recognition of different immunogenic patterns. Transcriptome profiling reveals an inventory of defense/stress-related genes that increase or acquire salt responsiveness after PRR elicitation and are characterized by overrepresentation of putative binding sites for WRKY transcription factors in their regulatory DNA sequences. Our genetic and biochemical studies suggest that PRR signaling utilizes WRKY40/WRKY18 to promote salt stress tolerance by accelerating salt-induced transcriptional reprogramming and by suppressing EDS1-based immunity. Recognition of non-pathogenic bacteria also leads to salt tolerance through these PRR signaling components. Our findings indicate that pattern sensing of cellular damage and plant-associated microbes is linked to salt stress tolerance via WRKY40/WRKY18-mediated transcriptional regulation.

## RESULTS

### Recognition of damage/microbe-associated molecular patterns leads to salt tolerance

Whole-genome microarray analysis for Pep2- and elf18-induced transcriptional reprogramming in *Arabidopsis* seedlings (Ross *et al*, 2014) produced an inventory of Pep2- and elf18-inducible genes (≧4-fold), i.e. 575 and 76 genes with Pep2 at 2 h and 10 h, respectively, and 536 and 380 genes with elf18 at 2 h and 10 h, respectively. *In-silico* data analysis suggests that the majority of these PTI-inducible genes are also induced in seedling shoots or roots in response to salt and osmotic stresses (Figure EV1A), as described previously for chitin (Espinoza *et al*, 2017). The common target genes included members of the *PROPEP* family and *PEPR1/PEPR2* (Figure EV1B), implying the extensive engagement of this DAMP pathway under salt stress. These data prompted us to examine whether recognition of different MAMPs and DAMPs leads to salt stress tolerance, and if so, by what mechanism.

We first tested whether pretreatment of seedlings with Pep, flg22 and elf18 peptides confers salt stress tolerance. Salt tolerance was determined as the ratio of viable (green) plants to dead/dying plants with bleaching leaves, over the total number of the tested plants. In non-elicited plants, the survival rate declined to 6% (175 mM NaCl), while survival of Pep1 pre-treated seedlings was 68%, 9 d after salt stress (Figure 1B). Pep1, Pep2, Pep3 and Pep4 pretreatments all significantly increased plant tolerance to 175 mM NaCl (Figures 1A-C, Table 1). PEPR1 recognizes all Pep peptides while PEPR2 recognizes only Pep1 and Pep2 (Krol *et al*, 2010; Bartels *et al*, 2013). Although it was previously described that PEPR1, but not PEPR2, is required for Pep3-triggered salt tolerance (Nakaminami *et al*, 2018), our analysis showed that Pep1-triggered salt tolerance was retained in *pepr1* or *pepr2* but abolished in *pepr1 pepr2* plants (Figures 1A, 1C, Table 1). This indicates that PEPR1 and PEPR2 both mediate salt tolerance, despite their differences in Pep ligand specificity.

**Figure 1.**
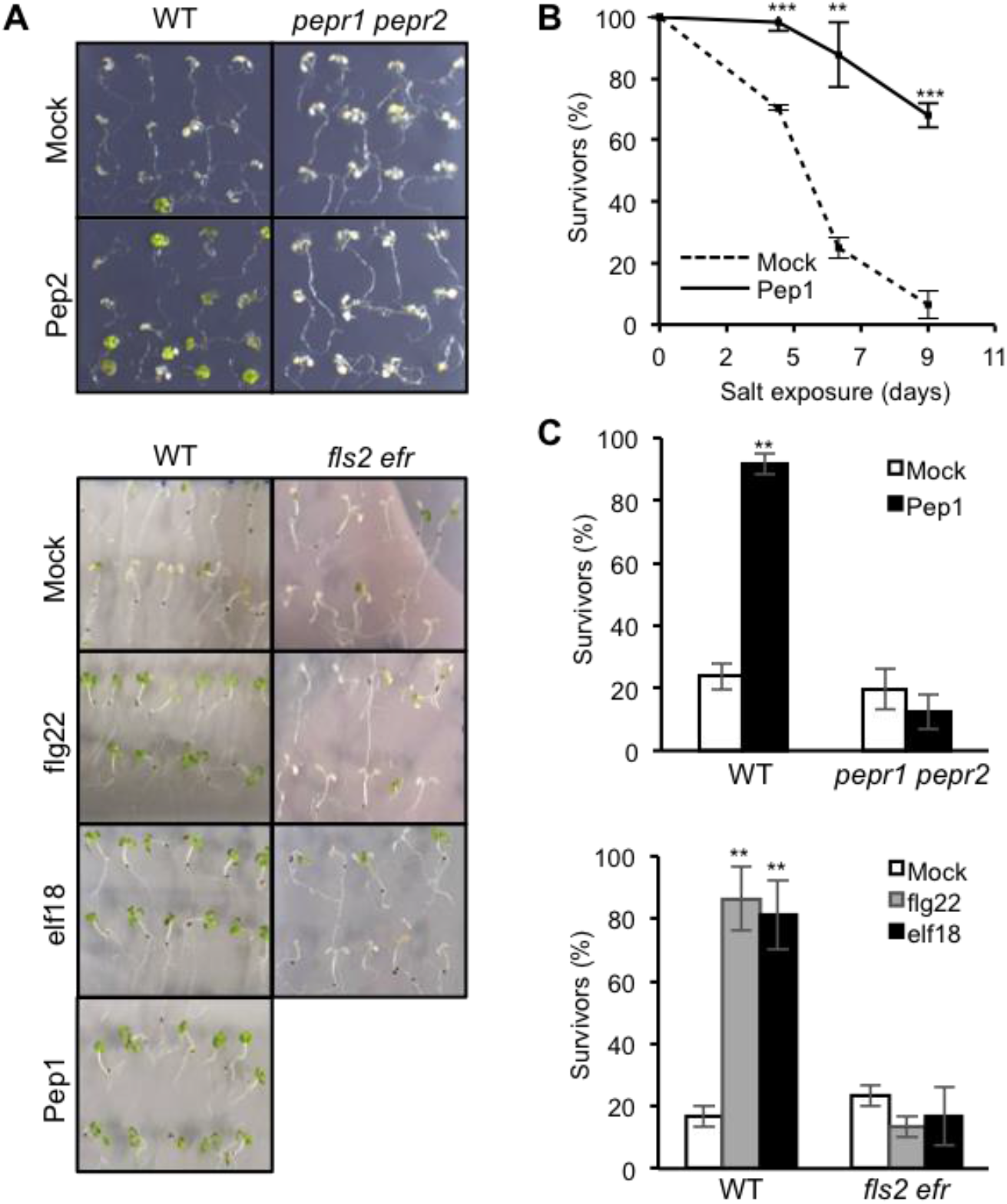
PRRs confers salt stress tolerance in *Arabidopsis thaliana* following recognition of cognate MAMPs/DAMPs. (**A**) Appearance of *A. thaliana* seedlings 6 days after exposure to 150 mM NaCl (Pep2) and ≥ 5 days after exposure to 175 mM NaCl (flg22, elf18 and Pep1) with or without the indicated MAMP/DAMP pretreatment. (**B**) Survival rate (mean ±s.e.m., n ≥30, 4 replicates) of WT seedlings after their exposure to 175 mM NaCl for the indicated times, with and without 0.1 μM Pep1 pretreatment. (***, *p* < 0.001 and **, *p* < 0.01 in two-tailed t-tests compared to the corresponding values of the mock-treated plants.) (**C**) Survival rate (mean ±s.e.m., n ≥ 20, 2 replicates) of *A. thaliana* seedlings 6 days after their exposure to 175 mM NaCl, with and without 0.1 μM the indicated MAMP/DAMP pretreatments. (**, *p* < 0.01 in Tukey’s honestly significant difference (HSD) tests compared to the value of mock-treated WT plants.)

**Table 1.** PEPR1 and PEPR2 both contribute to Pep-induced salt tolerance in *A. thaliana*. Survival rate (%) of seedlings 7 days after exposure to 175 mM NaCl.

| Genotype | Pretreatment | Survivors | Total seedlings | Survival rate (%) | Fisher's test (vs WT) | Fisher's test (vs mock) |
| --- | --- | --- | --- | --- | --- | --- |
| WT | Mock | 5 | 30 | 16.6 |  |  |
|  | Pep1 | 29 | 30 | 96.7 |  | P<0.01 |
|  | Pep3 | 30 | 46 | 65.2 |  | P<0.01 |
|  | Pep4 | 40 | 50 | 80.0 |  | P<0.01 |
|  | Mock | 6 | 136 | 4.4 |  |  |
|  | Pep2 | 87 | 140 | 62.1 |  | P<0.01 |
| *pepr1* | Mock | 5 | 35 | 14.3 |  |  |
|  | Pep1 | 26 | 30 | 86.7 | N.S. | P<0.01 |
| *pepr2* | Mock | 1 | 30 | 3.3 |  |  |
|  | Pep1 | 30 | 30 | 100 | N.S. | P<0.01 |

PRR signaling activation under sterile conditions typically leads to growth retardation (Boller & Felix, 2009). Conceivably, the lowered metabolic activity accompanying reduced growth could lower salt uptake into the plant, thereby conferring apparent tolerance. However, *pepr2* plants acquired salt tolerance following Pep1 application (Table 1), without discernible growth inhibition (Krol *et al*, 2010). Pep3 and Pep4 application also conferred salt tolerance without significantly inhibiting root growth (Figure EV1C; Table 1). This indicates that plant growth inhibition is not required for pattern-triggered salt tolerance, which we designate PTST.

Importantly, pretreatment with flg22 or elf18 also conferred salt tolerance through cognate PRRs (Figures 1A and 1C). These results indicate that PTST is not specific to an immunogenic pattern or receptor but is common to a broad range of MAMPs/DAMPs. This is consistent with the view established in plant immunity that a wide array of PRRs link the recognition of diverse cognate ligands to a largely overlapping set of defense outputs (Couto & Zipfel, 2016). The ligand dose dependence of flg22-induced salt tolerance was comparable with that of other flg22-induced outputs (Figure EV1D) (Gómez – Gómez *et al*, 1999; Aslam *et al*, 2009). These results suggest that PTST shares post-recognition signaling mechanisms with PTI across different PRR pathways. Notably, chitin application did not affect salt tolerance under our conditions, despite significant induction of a defense marker, *CYP71A13*, encoding cytochrome P450 involved in camalexin biosynthesis (Figures EV1E and EV1F).

### Pattern-triggered salt tolerance and immunity share early signaling steps downstream of the receptor

A major branch of PTI signaling triggered by the LRR-domain PRRs occurs through the receptor complexes with BAK1 and BIK1/PBL1 (Couto & Zipfel, 2016). To test possible *BAK1* dependence of PTST, we examined Pep1-triggered salt tolerance in the presence of a null *bak1-4* allele and a hypoactive *bak1-5* allele specifically impairing PRR-related BAK1 function (Roux *et al*, 2011; Schwessinger *et al*, 2011). Consistent with the previously described retention of PEPR-mediated defenses in a *bak1* null mutant (Yamada *et al*, 2016b), Pep1-triggered salt tolerance was unaffected in *bak1-4* (Figure 2). However, it was severely compromised in *bak1-5* plants and *bak1-5 bkk1* plants that additionally lack BAK1-related RK BKK1, as found for PEPR-mediated defenses (Yamada *et al*, 2016b)(Figure 2). There results show that PRR-regulating BAK1 function is required for PTST (Figure 2). Likewise, *bik1 pbl1* plants failed to acquire salt tolerance following Pep1 application (Figure 2), although they were not distinguishable from the wild type (WT) in flg22-induced salt tolerance (Figure 2). These results are consistent with PEPRs strictly requiring BIK1/PBL1 in PTI, while in contrast FLS2 does not (Zhang *et al*, 2010; Yamada *et al*, 2016a). Our results thus indicate that PTST signaling also occurs through these RLCKs.

**Figure 2.**
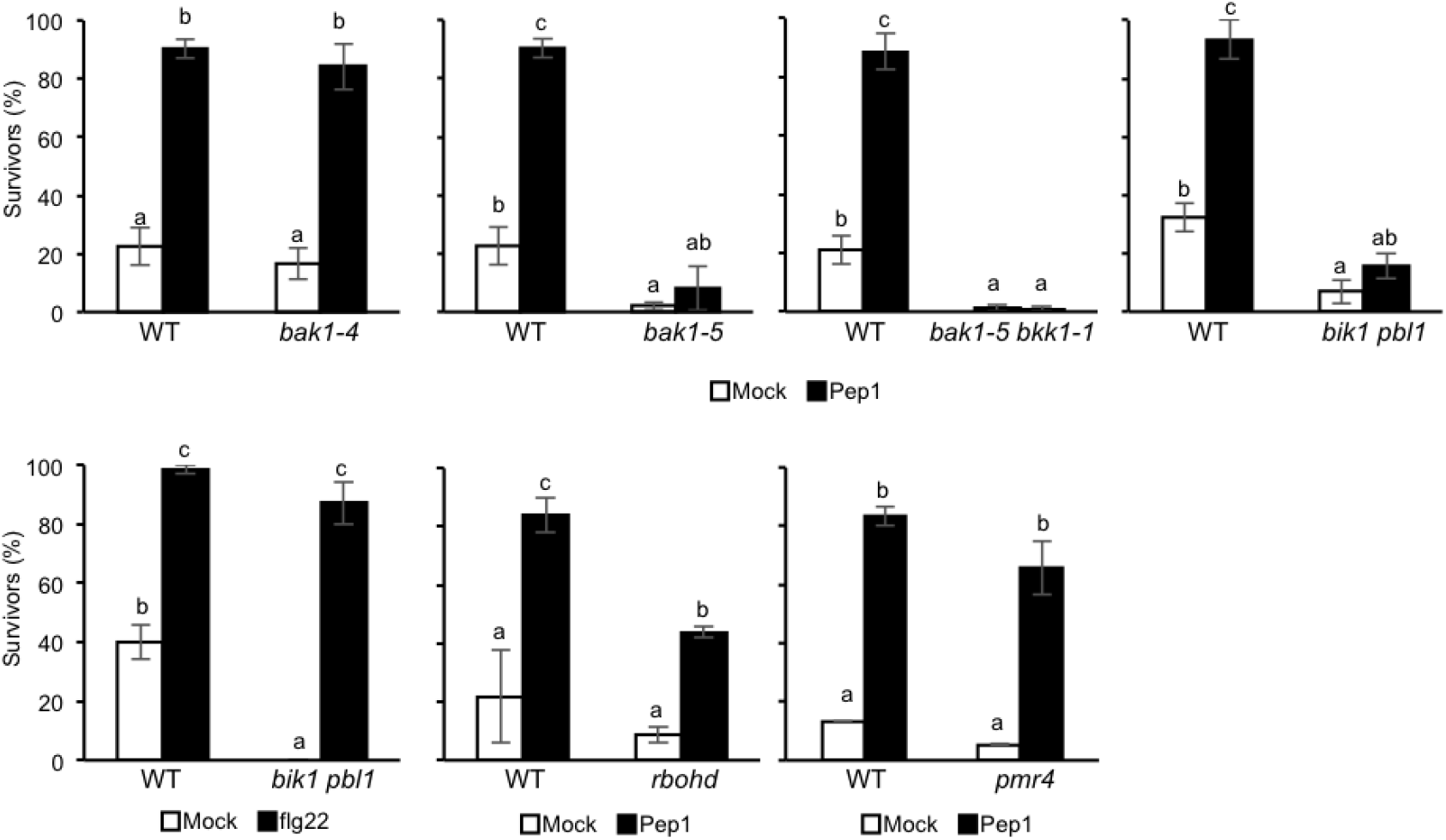
Genetic requirements for PTI signaling components in PTST. Following 0.1 μM Pep1/flg22 pretreatment, survival rate (mean ±s.e.m., 3 replicates unless otherwise stated) of seedlings was determined after their exposure to 175 mM NaCl for the indicated times. *bak1-4* and *bak1-5*, 9 days, n ≥ 20; *bak1-5 bkk1-1*, 8 days, n ≥ 30; *bik1 pbl1-Pep1*, 5 days, n ≥ 25; *bik1 pbl1*-flg22, 6 days, n ≥ 35, 2 replicates; *rbohd*, 8 days, n ≥ 30; *pmr4*, 6 days, n ≥ 30, 2 replicates. The letters above bars indicate p < 0.05 in Tukey’s HSD tests

Interestingly, seedling survival rate was also significantly lowered in *bak1-5, bak1-5 bkk1* and *bik1 pbl1* plants when directly exposed to salt stress without MAMP/DAMP application (Figures 2, mock controls), pointing to engagement of these PRR-associated kinases in salt tolerance. Our data suggest that the authentic receptor complexes mediate PTST and that DAMPs generated under salt stress signal via BAK1-dependent PRRs.

Pep1-triggered salt tolerance was reduced in *rbohd* plants lacking the PRR-associated NADPH oxidase responsible for a pattern-induced ROS burst (Figure 2; Kadota *et al*, 2015), pointing to a critical role also for this PRR output in PTST. By contrast, callose synthase PMR4/GSL5-dependent callose deposition (Kim *et al*, 2005) was not required for Pep1-triggered salt tolerance (Figure 2), demonstrating that PRR-induced callose deposition is dispensable for PTST. These data indicate that early signaling steps within and proximal to the PRR complexes, if not all, are shared with PTST.

### Pattern-triggered salt tolerance is robust against hormone perturbations

PRR signaling involves complex networks of defense-related hormones including SA, JA and ethylene in PTI (Pieterse *et al*, 2012). FLS2- and EFR-triggered immunity largely collapses in the simultaneous absence of *DDE2* encoding allene oxide synthase (AOS) required for JA biosynthesis, *EIN2* encoding the master regulator of ethylene signaling, *SID2* (*ICS1*) and *PAD4* (Tsuda *et al*, 2009). However, in *dde2 ein2 pad4 sid2* plants, Pep1-triggered salt tolerance was unaffected (Figure EV2A), indicating that these defense-related sectors are all dispensable for PTST.

We also assessed whether PTST is dependent on ABA, which is central to plant adaptation to salt, osmotic and water-deficit stresses (Cutler *et al*, 2010; Finkelstein, 2013). Pep-triggered salt tolerance was unaffected in *aba2-12* plants impaired in ABA biosynthesis (González-Guzmán *et al*, 2002) or in *areb1 areb2 abf3* plants lacking key transcription factors mediating ABA responses (Yoshida *et al*, 2015) (Figures EV2B and EV2C), suggesting that ABA is also dispensable for PTST. Overall, our findings point to high PTST robustness against perturbations of these biotic/abiotic stress-related hormone pathways.

### Damage sensing and signaling under salt stress involves the Pep-PEPR pathway

To test involvement of endogenous DAMPs in salt tolerance, we monitored endogenous PROPEP-PEPR signaling under salt stress. Given the substantial induction of *PROPEPs* and *PEPR1/PEPR2* in roots (Figure EV1A), we examined PROPEP3 protein expression in the roots of transgenic plants expressing *PROPEP3-Venus* under its native regulatory DNA sequences. A strong PROPEP3-Venus fluorescence signal was detected 24 h after salt stress, but not under non-saline conditions (Figure 3A). Damage-induced release of PROPEP1 from the vacuole and that of PROPEP3 to extracellular spaces (Hander *et al*, 2019; Yamada *et al*, 2016b) prompted us to test for possible PROPEP release under salt stress. We traced PROPEP3-Venus accumulation in an extracellular protein fraction recovered from the surrounding liquid medium, following salt stress and/or Pep1 application. Immunoblot analysis with the PROPEP3-specific antibodies (Ross *et al*, 2014) detected specific signals that were nearly of the predicted full-length size of PROPEP3-Venus (~10.4 + 27 kDa) following Pep1 application (Figure 3B), as described for Pep2 application (Yamada *et al*, 2016b). Apparently shorter forms of PROPEP3-Venus were detected under salt stress with or without Pep1 application (Figure 3B), possibly reflecting PROPEP3 processing that may occur in the intracellular or extracellular spaces. Under these conditions, endogenous PROPEP3-derived signals were not detected. Nevertheless, these results validate that PROPEP3 is released under salt stress.

**Figure 3.**
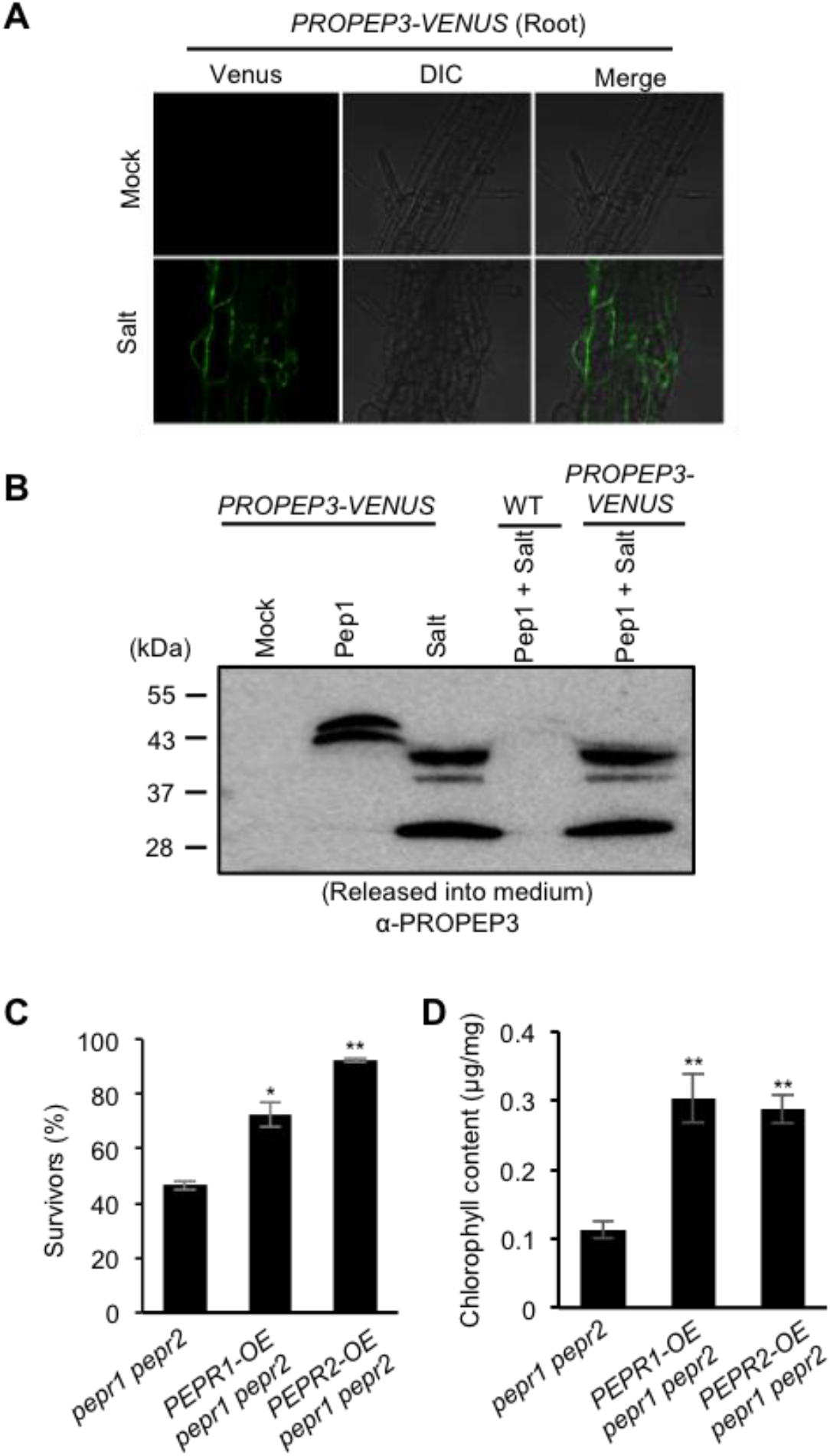
Endogenous PROPEP-PEPR signaling is activated under salt stress. (**A**) Live cell imaging of *promoter-PROPEP3::PROPEP3-VENUS* in *A. thaliana* roots under 150 mM NaCl for 24 h. (**B**) Immunoblot analysis for PROPEP3-VENUS in the extracellular fractions of seedlings when exposed to 0.5 μM Pep1, 150 mM NaCl or combinations thereof. Positions of the molecular weight markers shown on the left. Experiments were repeated twice with the same conclusions. (**C**) Survival rate (mean ± s.e.m, n ≥20, 2 replicates) of seedlings when directly exposed to 175 mM NaCl for 4 days, without MAMP/DAMP pretreatment. (**, p <0.01 and *p <0.05 in Tukey’s HSD compared to the WT values.) (**D**) Determination of chlorophyll contents (mean ±s.e.m., 4 replicates) in seedlings after their exposure to 750 mM sorbitol following pretreatement with 100 mM NaCl for 7 days. (**, p <0.01 in Tukey’s HSD tests and ***, p <0.001 in two-tailed t-tests compared to the values of *pepr1 pepr2* plants and WT plants, respectively.

To assess a possible contribution of the endogenous PEPR pathway to salt tolerance, we examined salt responses of PEPR1- and PEPR2-overexpressing lines (*PEPR1-OE* or *PEPR2-OE*, respectively) in the *pepr1 pepr2* background, without exogenous application of Peps or MAMPs. *PEPR1-OE* and *PEPR2-OE* plants both exhibited increased survival rates when directly exposed to 175 mM NaCl compared to that *of pepr1 pepr2* plants (Figure 3C). Moreover, following 1-week acclimatization to mild salt stress (100 mM NaCl), *PEPR1-OE* and *PEPR2-OE* plants acquired enhanced tolerance to severe osmotic stress (750 mM sorbitol) compared to *pepr1 pepr2* plants, indicated by the leaf chlorophyll contents (Figure 3D). These data provide compelling evidence that an endogenous PEPR pathway contributes to salt and osmotic stress tolerance, in the absence of exogenous Pep application. Collectively, the results indicate that salt stress-induced generation and release of PROPEP-derived peptides engages PEPR signaling in salt/osmotic stress tolerance.

### Pep1 pre-treatment strengthens transcriptome dynamics in response to salt stress

To gain a mechanistic insight into PTST, we performed transcriptome profiling in WT and *pepr1 pepr2* plants during the course of PTST. To capture useful information from the salt-sensitive samples, plants were subjected to 150 mM NaCl after Pep1 pretreatment. As salt-induced transcriptional reprogramming is largely achieved within the first 24 h (Geng *et al*, 2013), we obtained the data under salt stress for 3 and 24 h, after a 3-day Pep1 pretreatment (Figure 4A). Up- or down-regulated genes under salt stress in non-elicited plants (mock), with a cut-off of |log2 (fold change)| ≧ 1 (p <0.05), were assembled at the indicated times (Figure 4B), defining the salt-responsive differentially expressed genes (DEGs). Likewise, genes whose expression was significantly altered in salt responsiveness, both between Pep1- and mock-pretreated WT plants and between Pep1-pretreated WT plants and *pepr1 pepr2* plants, were assembled at the indicated times under salt stress (Figure 5B; Appendix Table 1), defining PTST-DEGs (exhibiting Pep1/PEPR-dependent alterations in salt responsiveness). DEGs were scored at the earliest time points when their expression levels first met these criteria.

**Figure 4.**
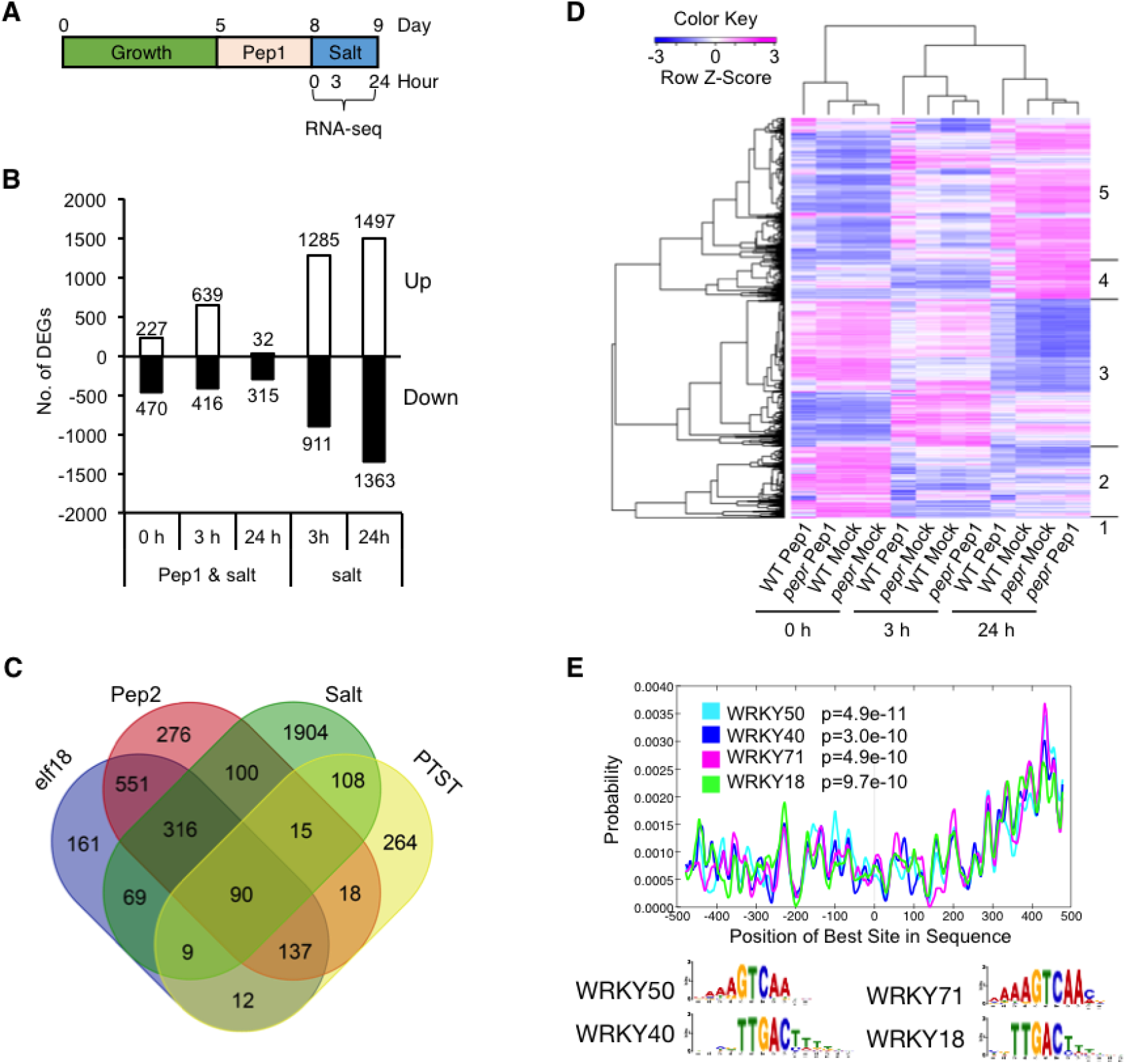
Rapid and strong activation of salt-induced transcriptional reprogramming during PTST. (**A**) Scheme for PTST settings subject to RNA sequencing analysis. (**B**) Numbers of Pep1- and/or salt-induced DEGs after exposure to 150 mM NaCl for the indicated times. (**C**) Venn diagram illustrating the overlap between elf18-, Pep2-, salt- and Pep1-PTST-inducible DEGs. Numerals represent the numbers of the genes. (**D**) A heatmap depicting salt-DEGs and Pep1-PTST DEGs after one minus Pearson correlation complete linkage hierarchical clustering. (**E**) *Cis*-element enrichment analysis with CentriMo for the regulatory DNA sequences within 1-kb (from 500 corresponding to the transcription starting sites to −500 on the X-axis) upstream of 343 genes in the Cluster 5, whose salt-induction was sensitized following Pep1 pretreatment. The results for the most over-represented 4 transcription factors are shown.

**Figure 5.**
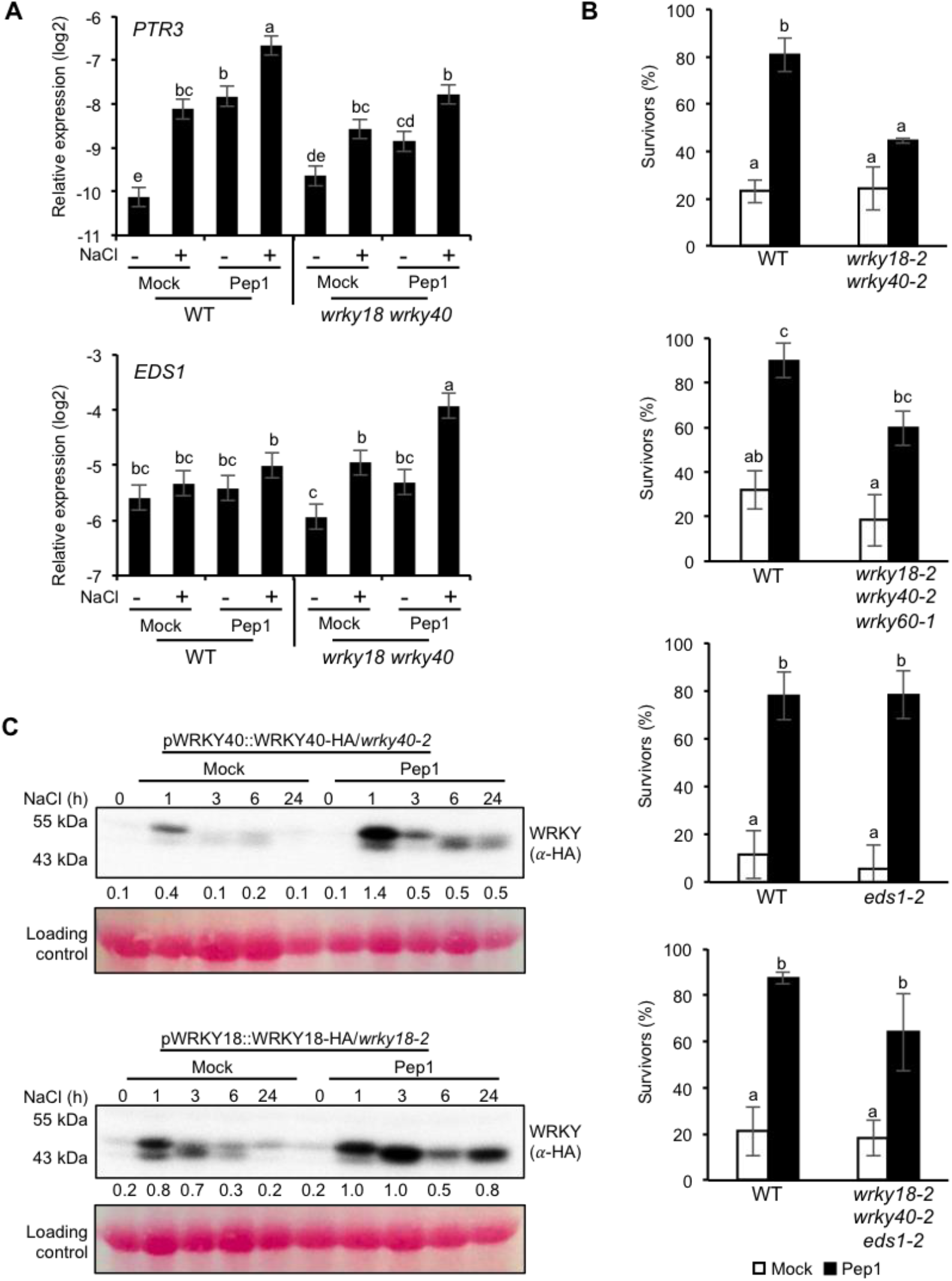
WRKY40 and WRKY18 transcription factors contribute to PTST in part by suppressing *EDS1*. (**A**) qRT–PCR analysis for *PTR3* and *EDS1* expression. Five-day-old seedlings were pretreated with 0.1 μM Pep1 for 4 days before exposure to 175 mM NaCl for 3 h. Data are mean ± s.e.m. of 3 biological replicates. Differences between samples were analyzed using a linear model, p < 0.01. (**B**) Survival rate (mean ± s.e.m.) of seedlings after exposure to 175 mM NaCl for 9 days, following 0.1 μM Pep1 pretreatment. Data represent mean ± s.e.m. of n ≥ 24, 3 replicates (*wrky18 wrky40*); n ≥ 30, 3 replicates (*wrky18 wrky40 wrky60*); n ≥ 25, 4 replicates (*eds1-2*); n ≥ 20, 5 replicates (*wrky18 wrky40 eds1-2*). The letters above bars indicate p < 0.05 in Tukey’s HSD tests. N.S., Not significant. (**C**) Immunoblot analysis for 9-day-old seedlings exposed to 175 mM NaCl for the indicated times following 0.1 μM Pep1 pretreatment. Positions of the molecular weight markers (left) and Ponceau S-stained loading controls (bottom) are shown. Experiments were repeated three times with the same conclusions. Numerals below the immunoblots indicate the band intensities relative to that of the corresponding loading control in the representative blots.

In non-elicited plants under salt stress, we detected a total of 1,285 up- vs. 911 down-regulated DEGs, and 1,497 up- vs. 1,363 down-regulated DEGs, at 3h and 24 h, respectively (Figure 5B). This suggests that salt-induced transcriptional reprogramming persisted over the tested time window. In Pep1-pretreated plants, we detected 639 up- vs. 416 down-regulated PTST-DEGs 3 h after salt stress, but merely 32 up- vs. 315 down-regulated PTST-DEGs at 24 h (Figures 4B and 4C). This suggests that PRR signaling particularly impacts the early responsiveness of salt-inducible genes.

Next, we examined possible overlap and divergence between the obtained salt-inducible DEGs and the previously described, Pep2- or elf18-responsive DEGs (2 h and 10 h; Ross *et al*, 2014). This showed that 599 genes (34.1% of Pep2/elf18-inducible genes and 22.9% of salt-inducible genes) were commonly induced between the two types of stimuli, while 1,155 and 2,012 genes were specifically induced in response to Pep2/elf18 and salt stress, respectively (Figure 4C). Our analysis indicates a substantial overlap, but also clear separation in the transcriptome between the biotic and abiotic stresses, in which a large portion of pattern-responsive genes is inherently not responsive to salt stress and *vice versa*.

Of 1,754 elf18- or Pep2-inducible DEGs and 2,611 salt-inducible DEGs, 281 genes (16.0%) and 222 genes (8.5%) were defined as PTST-DEGs, respectively (Figure 4C). Notably, these included pattern-specific DEGs which acquired salt inducibility following Pep1 pretreatment but were otherwise not responsive to salt stress: 3-d Pep1 treatment rendered 164 genes (125 + 39 genes in Figure EV3A, relative to 1,285 genes, inherently salt-inducible) significantly induced at 3 h, and 24 genes (13 + 11 genes in Figure EV3A, relative to 2,251 genes, inherently salt-inducible) at 24 h after salt stress. Moreover, PTST-DEGs included 264 genes which were not among the elf18/Pep2-DEGs or salt-DEGs, but acquired salt inducibility in Pep1-pretreated plants (Figure 4C). These results indicate that pre-activation of PRR signaling not only sensitizes salt stress responses but also broadens the range of target genes in salt stress responses, and emphasize that these effects are prominent early in salt responses.

### *WRKY18/WRKY40* transcription factors contribute to PTST

We further dissected all the salt- and PTST-DEGs (Figure 4C) by hierarchical clustering. The genes were classified into five clusters (Figure 4D, Appendix Table 1). Cluster 5 (2,194 genes) was over-represented with genes whose salt induction at 3h was increased after Pep1 pretreatment (Figure 4D). It included a set of genes related to both defense and salt stress responses. For example, *PTR3*, encoding a putative peptide transporter, promotes salt tolerance at seed germination and also basal resistance to *Pst* DC3000 (Karim *et al*, 2007, 2005). *SnRK2.8* encodes an osmotic stress-activated protein kinase, which promotes drought tolerance (Umezawa *et al*, 2014) and systemic immunity by phosphorylating NPR1 (Lee *et al*, 2015). Thus, it seems likely that PRR signaling pre-activation leads to faster establishment of a salt stress-adapted transcriptome during PTST.

To pursue this hypothesis, we assembled salt-inducible genes that exhibited faster induction following Pep1 pretreatment. Of the cluster 5 genes, 343 genes increased their salt induction at 3 h in Pep1-pretreated plants, with their induction higher at 24 h than 3 h in non-treated plants (Figure EV3B). In their regulatory DNA sequences, within 1000-bp upstream of the transcriptional start sites, a motif enrichment analysis (CentriMo, Bailey & Machanick, 2012) revealed over-representation of the W box-containing sequences (58 out of 59 over-represented transcription factor binding sites, Appendix Table 2). Four best-represented motifs were all prominent in the proximity to the transcription starting sites and included WRKY18- and WRKY40-specific DNA binding motifs (Figure 4E, Appendix Table 2), pointing to direct transcriptional regulation of these genes by WRKY18/WRKY40 during PTST. Interestingly, WRKY18/WRKY40 target genes (Birkenbihl *et al*, 2017a) were more clearly enriched in Cluster 5 genes displaying faster induction (149 out of 329 loci) compared to all Cluster 5 genes (471 out of 2083 loci) or PTST-DEGs (720 out of 5844 loci) (Figure EV3C), pointing to their role in rapid activation of a salt-induced transcriptome.

We next tested for involvement of the *WRKY18/WRKY40/WRKY60* subclade in PTST. Analyses of *wrky* mutant plants individually or simultaneously disrupted in the three *WRKY* members revealed that only *wrky18 wrky40* plants failed to develop PTST, in contrast to all the other tested mutants exhibiting salt tolerance, following Pep1 application (Figures 5B, Table 2). Similar PTST defects in an independent *wrky18 wrky40* mutant allele combination (Table 2), confirmed that *WRKY18/WRKY40* have an essential role in PTST. *WRKY60* is often functionally diverged from the other two *WRKY*s (Geilen & Böhmer, 2015; Shen *et al*, 2007). Accordingly, PTST was unaffected in *wrky18 wrky60* plants or *wrky40 wrky60* plants (Table 2), indicating that *WRKY60* is not required for PTST even in the absence of *WKRY40* or *WRKY18*.

**Table 2.**
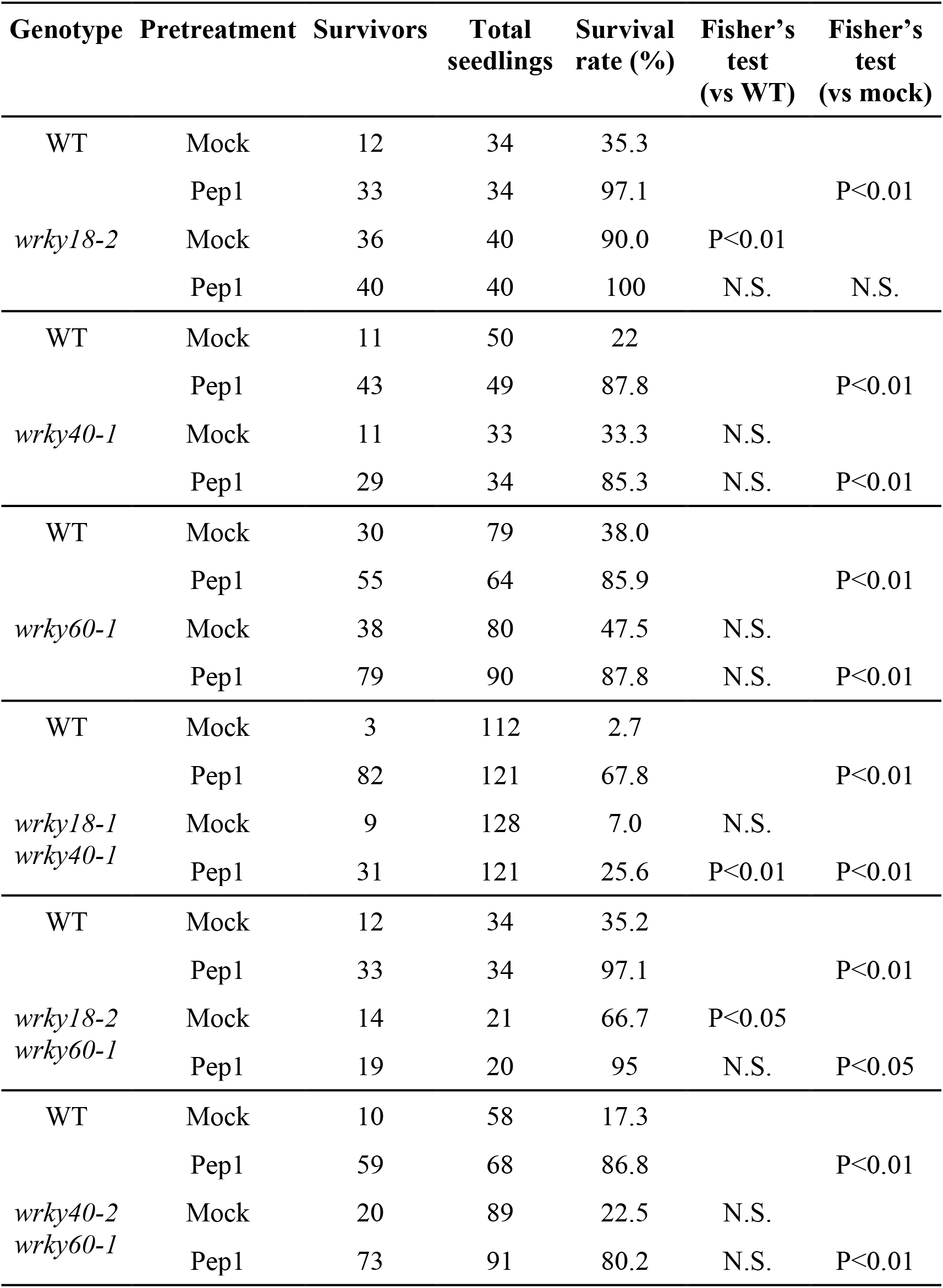
Roles for WRKY18/WRKY40/WRKY60 transcription factors in PTST. Survival rate after exposure to 175 mM NaCl for 7 days (or 9 days for *wrky18-1 wrky40-1*) following 0.1 μM Pep1 pretreatment. N.S., Not significant; **, p <0.01 in Tukey’s HSD tests.

Importantly, basal salt tolerance was enhanced in non-elicited (mock-treated) *wrky18* plants and *wrky18 wrky60* plants, but was lowered to WT-like levels in *wrky18 wrky40* plants (Figures 5B and Table 2). This suggests that *WRKY18* and *WRKY40*, respectively, negatively and positively influence basal salt tolerance, and that *WRKY40* confers salt tolerance in the absence of WRKY18, while *WRKY60* has a minor role. Remarkably, *WRKY18* contributed to PTST in the absence of *WRKY40* (see *wrky40* plants vs. *wrky18 wrky40* plants in Figure 5B and Table 2), implying that PRR signaling, and possibly also the loss of *WRKY40*, switches *WRKY18* from a negative to positive regulator of salt tolerance. To our surprise, however, the loss of *WRKY60* largely restored PTST to *wrky18 wrky40* plants, as shown in *wrky18 wrky40 wrky60* plants (Figure 5B), suggesting that *WRKY60* negatively influences PTST in the absence of *WRKY18* and *WRKY40*. Our findings uncover a balanced, compensatory *WRKY* transcription factor network in which *WRKY18* covers for loss of *WRKY40* in PTST, while there is antagonism between *WRKY18* and *WRKY40* in basal salt tolerance, and between *WRKY18*/*WRKY40* and *WRKY60* in PTST.

### Sustained WRKY18/WRKY40 accumulation during PTST for promotion of salt tolerance and repression of immunity regulator EDS1

To gain a mechanistic insight into the *WRKY40/WRKY18* function, we examined some of their target genes during PTST. Direct target genes of WRKY18/WRKY40 during PTI includes *PTR3* (Birkenbihl *et al*, 2017a). *PTR3* was induced in response to Pep1 or salt stress alone, and its salt induction was enhanced at 3 h as well as at 24 h in Pep1-pretreated plants (Figures 5A and EV3D). In *wrky18 wrky40* plants, however, Pep1 stimulation of *PTR3* expression was lowered with or without salt stress, although its salt inducibility in non-treated plants (mock) was unaffected (Figure 5A). These data are consistent with a specific impairment of PTST, but not basal salt tolerance, in *wrky18 wrky40* plants, and with the idea that positive regulators of salt tolerance are among the transcriptional targets of WRKY18/WRKY40 during PTST.

*EDS1* is another direct target of WRKY18/WRKY40-mediated transcriptional repression in pathogen defense (Birkenbihl *et al*, 2017a; Pandey *et al*, 2010). *EDS1* expression was largely unaltered in response to Pep1, salt stress or their combination, but was significantly increased in *wrky18 wrky40* plants under salt stress with or without Pep1 pretreatment (Figures 5A and EV3D). Given that EDS1-mediated defense activation disables salt-induced acquisition of osmotic tolerance (Ariga *et al*, 2017), we tested for a possible functional relationship between *WRKY18/WRKY40* and *EDS1* in PTST. Strikingly, PTST defects in *wrky18 wrky40* plants were largely rescued in *eds1 wrky18 wrky40* plants (Figure 5B), indicating that in the absence of *WRKY18/WRKY40 EDS1* de-repression causes PTST dysfunction. In the presence of *WRKY18/WRKY40* however, the loss of *EDS1* or *PAD4* alone did not significantly affect basal or Pep1-induced salt tolerance (Figures 5B and EV2A). These phenotypes, put together with WT-like basal salt tolerance in *wrky18 wrky40* plants, suggest that *WRKY18/WRKY40*-mediated suppression of *EDS1* function is specifically required for PTST, as *EDS1* de-repression could boost PRR-triggered signaling towards detrimental defense activation. Hence, our findings identify a critical role for dual WRKY18/WRKY40 functions during PTST: the transcriptional activation and repression of positive (eg. *PTR3*) and negative (eg. *EDS1*) regulators of salt tolerance, respectively, when PRR signaling is activated.

We examined how PRR signaling involves WRKY18 and WRKY40 in PTST. To this end, we conducted immunoblot analyses of functional HA-tagged WRKY18 and WRKY40 proteins expressed under the control of their native regulatory DNA sequences (*pWRKY18::WRKY18-HA wrky18* and *pWRKY40::WRKY40-HA wrky40*, respectively; Birkenbihl *et al*, 2017a). WRKY18 and WRKY40 accumulation was shown to be rapidly induced in response to flg22, with a peak of protein abundance at 1.5 h (Birkenbihl *et al*, 2017a). WRKY18/WRKY40 accumulation was reduced to nearly background levels 4 d after Pep1 application (0 h NaCl, Figure 5C). WRKY40-HA accumulation became strongly induced 1 h after salt stress, and then diminished (Figure 5C), indicating there is transient WRKY40 induction during PTI and salt stress. Importantly, Pep1 pretreatment markedly elevated and prolonged salt-induced WRKY40-HA accumulation up to 24 h (Figure 5C), following its increased mRNA expression (Figure EV3D). A similar Pep1 effect was observed for WRKY18-HA accumulation (Figure 5C). These results show that PRR signaling pre-activation leads to enhanced and durable accumulation of both WRKY40 and WRKY18 under salt stress.

### Non-pathogenic bacteria confer PTST

Having shown that bacterial MAMP application confers salt tolerance (Figure 1C), we tested whether immune recognition of bacteria also leads to salt tolerance. To this end, we determined the effects of pre-inoculation with different strains of *Pst* DC3000 on salt stress tolerance: *Pst* DC3000 *ΔhrpS*, impaired in the expression of the type III effectors (Hutcheson *et al*, 2001) and conventionally used as a PTI trigger, and *Pst* DC3000 *AvrRpm1* or *Pst* DC3000 *AvrRps4*, inducing ETI conferred by the CC-NLR RPM1 and the TIR-NLR pair RRS1-S/RPS4, respectively (Grant *et al*, 1995; Gassmann *et al*, 1999; Saucet *et al*, 2015). All of these bacterial strains fail to grow in the WT plants used here which harbor the cognate NLRs. Pre-inoculation with *Pst* DC3000 *ΔhrpS* significantly enhanced the survival rate of seedlings under salt stress, whereas *Pst* DC3000 *AvrRpm1* or *Pst* DC3000 *AvrRps4* did not (Figure 6A). Without salt stress, plant survival rates were essentially indistinguishable between these non-pathogenic and avirulent strains (Table EV1). These results suggest that PRR recognition, but not NLR recognition of live bacteria, effectively confers salt tolerance.

**Figure 6.**
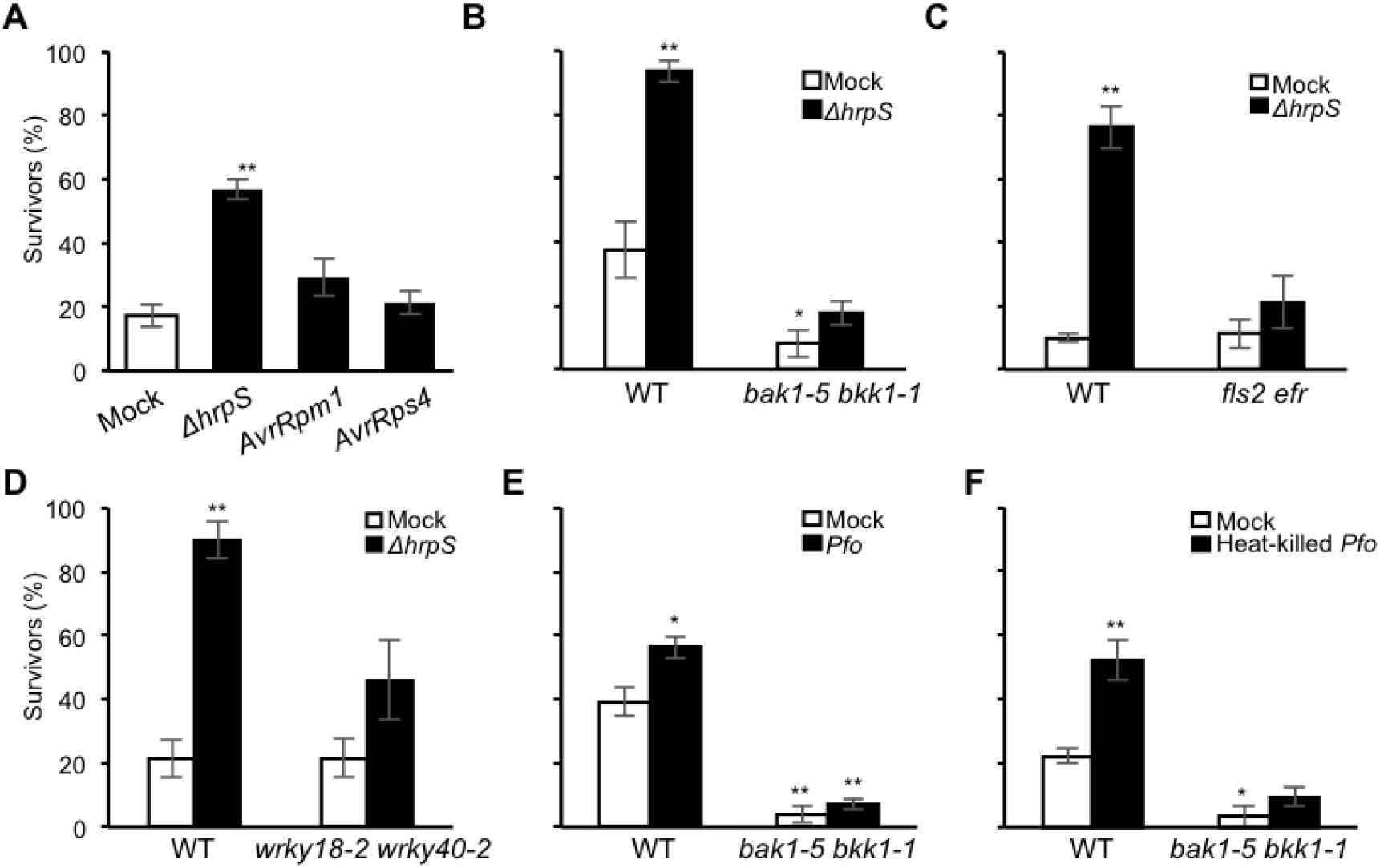
Non-pathogenic bacteria confer salt tolerance through the host PRR-WRKY40/WRKY18 module. (**A**) Survival rate (mean ± s.e.m., n ≥ 25, 3 replicates) of WT seedlings after exposure to 175 mM NaCl for 5 days following transient inoculation with the indicated *Pst* DC3000 strains. (**, p <0.01 in Tukey’s HSD tests compared to the values of the mock control.) (**B**-**F**) Survival rate (mean ± s.e.m., n ≥ 20, 3 replicates in B, D, E and F or 4 replicates in C) of seedlings exposed to 175 mM NaCl for 5 days (B, C, E), 6 days (D) and 4 days (F) following inoculation with the indicated *Pst* DC3000 strains. (**, p < 0.01 and *, p < 0.05 in Tukey’s HSD tests compared to the values of mock-treated WT plants.)

Notably, bacterium-triggered salt tolerance was strongly reduced in the *bak1-5 bkk1-1* and *fls2 efr* mutants (Figures 6B-C), as in MAMP/DAMP-triggered salt tolerance (Figure 1, Figure 2). Basal salt tolerance under sterile conditions (in mock controls without bacteria) was significantly lowered in *bak1 bkk1* plants but was unaffected in *fls2 efr* plants (Figures 6B and C), pointing to involvement of a BAK1-dependent DAMP receptor(s) but not MAMP receptors FLS2/EFR in basal salt tolerance. Moreover, bacterium-triggered but not basal salt tolerance was compromised in *wrky18 wrky40* plants (Figure 6D), again pointing to a specific contribution of *WRKY18/WRKY40* to PTST but not to basal salt tolerance. These results indicate that the PTST response module of PRRs and WRKY18/WRKY40 becomes engaged in response to bacterial challenge, thereby conferring salt tolerance. Assays with another non-pathogenic PTI-triggering bacterium, *Pseudomonas simiae* WCS417 (*Pfo* abbreviated after *Pseudomonas fluorescence*) also conferred salt tolerance which was abolished in *bak1-5 bkk1* plants (Figure 6E), strengthening the view that bacteria confer salt tolerance through BAK1-dependent PRR signaling.

Finally, we tested whether bacterial MAMP recognition without live bacteria is sufficient to acquire salt tolerance. Indeed, inoculation with heat-killed *Pfo* enhanced salt tolerance in a *BAK1/BKK1*-dependent manner (Figure 6F). These results suggest that PRRs are important for salt stress sensing and adaptation when recognizing molecular patterns derived from the host-associated microbes or cellular damage.

## DISCUSSION

Immune receptor activation can positively or negatively influence abiotic stress responses, yet the molecular logic behind this signaling plasticity remains poorly understood. Here, we show that PRR signaling triggers an enhanced or primed state of salt stress tolerance in plants (Figure 1, Table 1). Several lines of evidence indicate that PTST and PTI share previously described key steps within and proximal to the receptor complexes, at least for three BAK1-dependent PRR pathways. A failure to mount PTST in the *bak1-5* mutant and in the absence of *BIK1/PBL1or RBOHD* indicates that PTST is achieved by authentic PRR signaling (Figure 2). Effective cross-tolerance to biotic and salt stresses following PRR signaling may reflect similar cellular states and requirements in these stress conditions. This notion is supported by a substantial overlap between the pattern-induced and salt-induced transcriptomes (Figures 4 and EV1). Consistently, pattern recognition leads to the sensitization of salt-responsive genes and mobilization of otherwise non-responsive genes, most prominently during early responses to salt stress (Figures 4, EV1 and EV3). These findings indicate rapid activation and expansion of the salt-responsive transcriptome as an important basis for PTST, and predict the existence of an intermediate critical step in PRR signaling for this output.

By focusing on genes whose salt induction is strengthened and/or accelerated following Pep1 application, we revealed an interesting set of PTST-characteristic DEGs (Figure 4D, Appendix Table 1). A large portion of these genes seem to to be directly regulated by WRKY18/WRKY40, as WRKY18/WRKY40 DNA-binding sites are strongly over-represented in their regulatory DNA sequences (Figure 4E, Appendix Table 2). Moreover, a specific *WRKY18/WRKY40* requirement in PTST suggests that PRR signaling engages these two transcription factors to facilitate rapid mobilization of salt-adaptive pathways during PTST. The inventory obtained here for the PTST target genes implies two functional roles for WRKY18/WRKY40 in transcriptional regulation during PTST: Up-regulation of positive regulators and down-regulation of negative regulators of salt tolerance. In addition to *PTR3*, *WRKY18* and *WRKY40* themselves undergo positive transcriptional regulation (Figures 5A, EV3D), which leads to increased and sustained accumulation of the two WRKY proteins under salt stress. Because this boost is maintained after pattern-induced mRNA/protein expression is diminished (Figure 5C), *WRKY18/WRKY40* expression is primed as a critical output of PRR signaling during PTST. On the other hand, following PRR signaling, *WRKY18/WRKY40* ensures that induction of *EDS1* is prevented under salt stress (Figure 5A), which is detrimental to salt/osmotic stress tolerance (Ariga *et al*, 2017). Our findings thus suggest that limiting the magnitude of immune activation is a key for linking PRR-triggered signaling to salt tolerance (Figure 7). They also reconcile the apparent discrepancy between positive and negative effects of immune activation on salt/osmotic tolerance (Saijo & Loo, 2020; this study), and suggest that the EDS1 activation state is a critical decision node between abiotic-biotic stress cross-tolerance and tradeoff.

**Figure 7.**
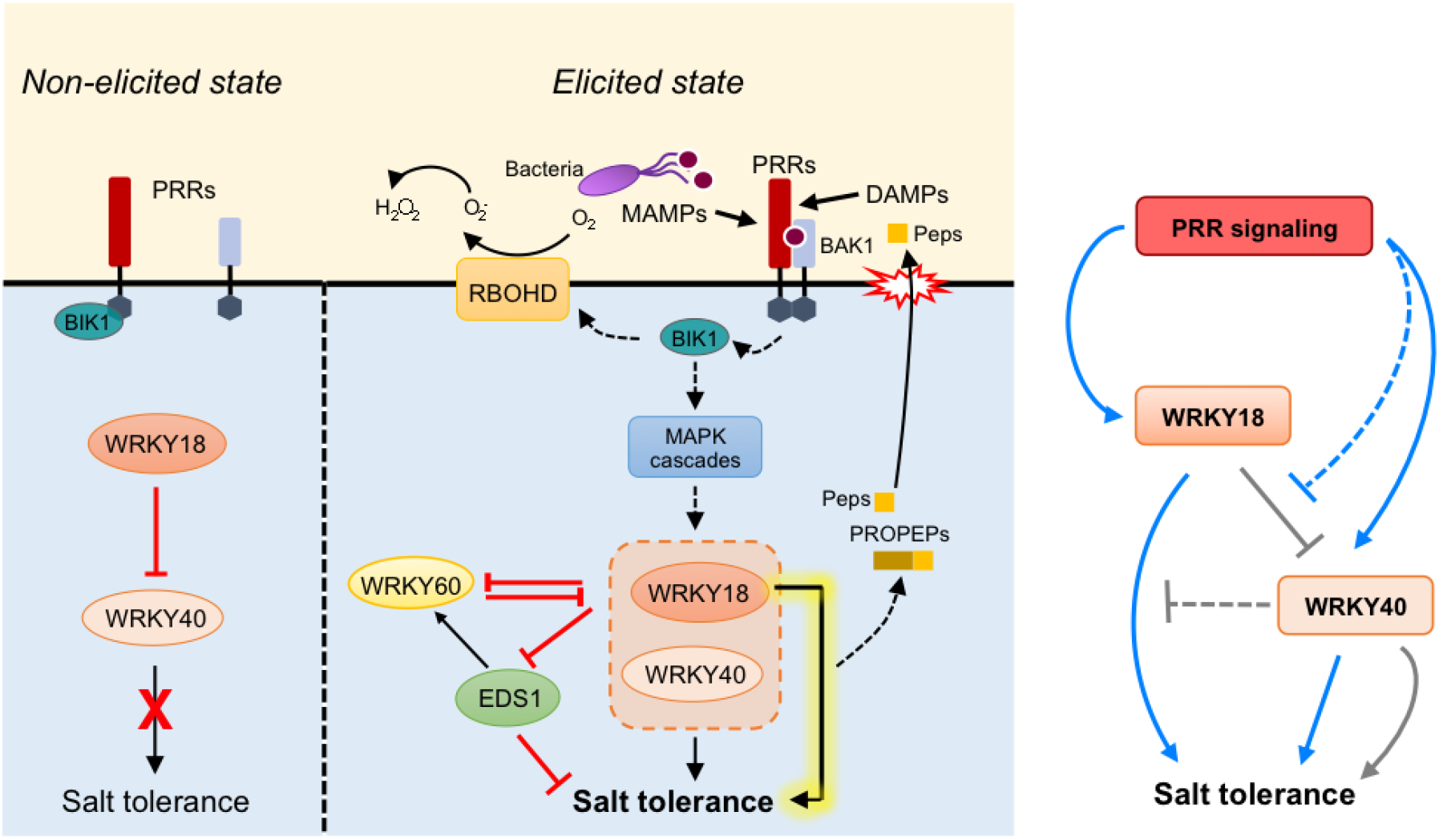
A model for PRR signaling and incoherent feed-forward loop consisting of WRKY18/WRKY40 during PTST. Following recognition of cognate MAMP/DAMP ligands, PRRs trigger signaling cascades through the previously described PRR complexes and signaling regulators, which result in priming of *WRKY40/WRKY18* expression. This leads to enhanced and sustained accumulation of WRKY40/WRKY18 under salt stress, a possible basis for rapid activation of the salt-adaptive transcriptome during PTST, including up-regulation of salt tolerance executors and down-regulation of immunity activators such as *EDS1*. Our findings point to a model in which salt stress stimulates presentation of DAMPs from cellular damage and MAMPs from plant-associated microbes, thereby involving PRRs in salt stress sensing and signaling that is linked via WRKY40/WRKY18 to salt tolerance. An incoherent type 4 feed-forward loop illustrating *WRKY18-WRKY40* functional interactions is shown in the right. PRR signaling leads to prioritization of *WRKY18* promotion of salt tolerance over its suppression of *WRKY40*-mediated salt tolerance. Dotted lines indicate the actions/links hypothesized.

Functional interactions between WRKY18 and WRKY40 play an important role in balancing immunity and salt tolerance following PRR activation. In PTST, *WRKY40* acts as a primary positive regulator of salt tolerance and its role is supported by *WRKY18*, while *WRKY40*-mediated salt tolerance is suppressed by *WRKY18* in the absence of PRR pre-activation (Figure 5B, Table 2). Plasticity of *WRKY18*-*WRKY40* interactions conforms to an incoherent type-4 feed-forward loop (I4-FFL; Alon, 2007), in which *WRKY18* (a regulator) represses *WRKY40* function (another regulator) that positively regulates salt tolerance (the target), while *WRKY18* alone has a positive influence on salt tolerance (illustrated in Figure 7). In non-elicited plants, *WRKY18* repression of *WRKY40*-mediated salt tolerance dominates, explaining the enhanced and WT-like salt tolerance in *wrky18* and *wrky18 wrky40* plants, respectively (Figure 5B, Table 2). PRR signaling utilizes *WRKY18* and *WRKY40* redundantly to promote salt tolerance during PTST, indicated by the retention and impairment of PTST, respectively, in the *wrky* single and *wrky18 wrky40* double mutants (Figure 5B, Table 2). Functional targets of the two WRKYs in PTST include suppression of *EDS1* and *WRKY60*, indicated by PTST restoration in *wrky18 wrky40 eds1* and *wrky18 wrky40 wrky60* mutants (Figure 5B). *WRKY40* also functions with the transcription factor *BZR1* in suppression of PTI via the phytohormone brassinosteroid (Lozano-Durán *et al*, 2013). Hence, *WRKY18* repression of *WRKY40* likely serves to restrict the attenuation of immunity. Conversely, PRR signaling switches *WRKY18* from a negative to a positive regulator that reinforces *WRKY40*-mediated immune attenuation and salt tolerance, thereby increasing robustness of PTST. A similar I4-FFL has been described in PTI, in which JA signaling provides robust and tunable control of SA-based defenses (Mine *et al*, 2017). Balancing different cellular demands that often mutually conflict (eg. biotic vs abiotic stress responses or stress tolerance vs plant growth) is crucial for plants to adapt to fluctuating environments. The biological significance and possible prevalence of signaling regulation via I4-FFLs merits further investigation under combined stress conditions.

Importantly, *WRKY18/WRKY40* function is impacted by *WRKY60*. In the absence of *WRKY40/WRKY18*, *WRKY60* suppresses PTST (Figure 5B). In this respect, we note that *WRKY60* is induced during *EDS1*-dependent transcriptional reprogramming in basal immunity (Cui *et al*, 2017), implying that *EDS1* suppresses PTST in part through *WRKY60*-*WRKY40/WRKY18* antagonism. Although *WRKY18* works together with *WRKY40* and *WRKY60* to attenuate basal resistance against *Pst*, gain of *WRKY18* function alone enhances bacterial resistance (Xu *et al*, 2006). Also, *WRKY18* alone promotes systemic acquired resistance (Wang *et al*, 2006). Functional relationships among the three *WRKY* members also change in ABA signaling. They act as negative regulators of ABA signaling in seed germination and seedling growth, with a primary and secondary role provided by *WRKY40* and *WRKY18*, respectively (Yan *et al*, 2013; Shang *et al*, 2010), while *WRKY40* and *WRKY18/WRKY60*, respectively, display negative and positive effects on ABA responsiveness in a different stress context (Chen *et al*, 2010). The molecular mechanisms underlying intricate cross-regulation of *WRKY18/WRKY40/WRKY60* remain obscure. WRKY18, WRKY40 and WRKY60 physically associate with themselves and each other to alter their DNA binding activities: WRKY40 *in vitro* binding to W-box containing DNA sequences is increased or lowered, respectively, by the presence of WRKY18 or WRKY60 (Xu *et al*, 2006). WRKY18 and WRKY40, but not WRKY60, localize within nuclear bodies containing the red/far-red light receptor Phytochrome B, under non-saline and non-elicited conditions (Geilen & Böhmer, 2015). WRKY40 and WRKY60, but not WRKY18, are refractory to *Pst* effector PopP2-mediated acetylation and disruption of DNA binding (Le Roux *et al*, 2015). Future studies are required to elucidate how the properties of these three WRKY proteins relate to their specific functions and cross-regulation.

DAMPs represent a common signature of biotic and abiotic stress conditions in animals and plants (Gust *et al*, 2017; De Lorenzo *et al*, 2018). In plants, abiotic modulation of cell walls and phospholipid membranes generates a battery of DAMPs (Chen *et al*, 2020; Rui & Dinneny, 2020; Herger *et al*, 2019; Jiang *et al*, 2019). Although the identity of cognate DAMP ligands remains elusive, different RKs are involved in mediating PTI-like defense responses and salt tolerance under salt stress conditions (Feng *et al*, 2018; Engelsdorf *et al*, 2018; van der Does *et al*, 2017). Here, we show that PROPEP3 and their-derived short fragments are released following salt stress, without microbes or exogenous MAMP/DAMP application. *PROPEP2/PROPEP3* expression represents an important preparatory step for positive feedback of defense signaling through PEPRs (Ross *et al*, 2014). *PROPRP2/PROPEP3* were among the 343 genes displaying faster salt induction following PRR activation (Figure EV3D), pointing to a role for the PEPR pathway in rapid mobilization of salt-adaptive responses during PTST. Indeed, *PEPRs* limit salt tolerance and salt-induced osmotic stress tolerance, both under sterile conditions (Figures 3C and 3D). Genetic requirements for *BAK1* and *BIK1/PBL1* (Figure 2) are consistent with the involvement of BAK1/BIK1-dependent DAMP receptors, including PEPRs, in salt tolerance. These findings strengthen the view that PRRs contribute to salt tolerance.

Shared use of common signaling components between PTI and salt tolerance extends beyond BAK1/BIK1-dependent PRR pathways. Glycosyl inositol phosphorylceramide sphingolipids provide Na^+^ sensors to induce Ca^2+^ influx for SOS signaling under salt stress (Jiang *et al*, 2019), and also perception sites for bacterial/fungal/oomycete Necrosis and ethylene-inducing peptide 1-like (NLP) proteins (Lenarčič *et al*, 2017). Salt tolerance is limited by the *Catharanthus roseus* RK *FER* (Feng *et al*, 2018; Zhao *et al*, 2018). FER recognizes immuno-stimulatory and immuno-suppressive members of the endogenous RALF peptides and also scaffolds different PRR complexes (Stegmann *et al*, 2017; Haruta *et al*, 2014). FER-mediated salt tolerance in part depends on its ability to bind pectin and protect pectin crosslinking, suggesting its role in the sensing and management of cell wall integrity under salt stress (Feng *et al*, 2018). Following S1P subtilase cleavage, RALF22/RALF23 are released from LRR-containing extensins LRX3/LRX4/LRX5, thereby lowering salt tolerance through FER (Zhao *et al*, 2018). Notably, S1P-cleaved RALF members attenuate both FER-mediated salt tolerance and PTI (Zhao *et al*, 2018; Stegmann *et al*, 2017). These studies further highlight the resemblance of PTI and salt stress signaling. Under our conditions, however, chitin signaling pre-activation failed to confer salt tolerance. The apparent discrepancy between our and previous studies of chitin/CERK1-mediated salt tolerance (Figure EV1; Brotman *et al*, 2012; Espinoza *et al*, 2017) might reflect a divergence between different ectodomain classes of PRRs in their optimal conditions for salt tolerance, as seen in their regulation of immunity (Saijo *et al*, 2018).

Successful induction of PTST by PRR recognition of bacterial MAMPs, but not by NLR recognition of their effectors (Figures 1 and 6), fits with the idea that strong activation of immunity negatively influences salt tolerance. This predicts the existence of a critical threshold beyond which further immune activation comes at a cost for salt stress tolerance. Recent studies show that PRR signaling provides an integrating basis for ETI, and that mutual PTI-ETI potentiation is required for effective pathogen resistance (Ngou *et al*, 2020; Yuan *et al*, 2020). At present, how NLR signaling exceeds the predicted threshold during ETI remains poorly understood. In barley, the CC-NLR MLA was reported to confer ETI against powdery mildew fungi in part by removing immune-suppressive functions of *HvWRKY1* and *HvWRKY2*, which are orthologous to *Arabidopsis WRKY18/WRKY40/WRKY60* (Shen *et al*, 2007; Liu *et al*, 2014). This suggests a conserved role for this *WRKY* subclade in modulating immunity thresholds.

## MATERIALS AND METHODS

### Plant materials and growth conditions

The *Arabidopsis thaliana* accession Col-0 was used as WT. Plant materials used are provided in Appendix Table 3. Seeds were sterilized with 6% sodium hypochlorite and 0.1% Triton X-10 for 15 minutes, rinsed 5 times with autoclaved distilled water and stratified at 4 °C for 2-5 days before use. The growth medium used was Skoog-Murashige (MS) medium (1/2 strength MS basal salts, 25 mM sucrose, 0.5 g/L MES, pH 5.7) unless otherwise stated. Plants were grown under 14 h light/ 10 h dark at 22 °C unless otherwise stated. For detection of extracellular PROPEP3-Venus protein, two-week-old seedlings in liquid growth media were exposed to 0.5 μM Pep1 for 3 days, 150 mM NaCl for 3 days or 0.5 μM Pep1 for 12 h followed by 150 mM NaCl for 3 days under standard growth conditions.

### Pattern-triggered salt tolerance assay

Four-day-old seedlings in the liquid growth media were treated with the indicated elicitors (0.1 μM Peps/flg22/elf18, 100 μg/ml chitin). For treatment with heat-killed bacteria, bacteria cultivated (as descried below) up to OD_590_ 0.2 were collected, suspended and then autoclaved at 121°C for 20 minutes. The supernatants after centrifugation were recovered for use. Four days after elicitor/bacterium treatments, seedlings were transferred to the agar growth media supplemented with 175 mM NaCl. The number of viable seedlings was scored every day for the indicated duration. Survival ratio was determined as the number of viable seedlings relative to the total number of seedlings used.

### Acquired osmotic tolerance assay

Assays for salt-induced osmotic stress tolerance were performed as described in Ariga *et al*, 2017. In brief, seven-day-old seedlings were transferred from agar growth media to that supplemented with 100 mM NaCl, and further incubated for 7 days. Seedlings were then transferred to that supplemented with 750 mM sorbitol, and grown for another 14 days before the determination of chlorophyll contents.

### Quantitative RT-PCR analysis

Total RNA was extracted from plant samples with Purelink (Nacalai Tesque, Japan) and reverse transcribed with PrimeScript Reagent Kit Perfect Real Time (Takara, Japan) according to manufacturer’s instructions. qRT-PCR was performed with Power SYBR Green PCR Master Mix (Applied Biosystems, Japan) using the Thermal Cycler Dice RealTime TP870 (Takara, Japan) under the following conditions: 50°C 2 min, 95°C 10 min, 95°C 15 s followed by 60°C 1 min for 40 cycles, then 95°C 15 s, 60°C 30 s, and finally 95°C 15 s. The primers used are provided in Appendix Table 4.

### Protein extraction and immunoblot analysis

Protein extracts were prepared by homogenizing frozen tissues in a lysis buffer [50 mM Tris-HCl pH7.5, 2% SDS, 2mM DTT, 2.5 mM NaF, P9599 protease inhibitor cocktail (Sigma)] for 15 min at room temperature. The supernatants recovered after centrifugation at 13,000 g for 15 minutes were subjected to immunoblot analysis on 10% SDS-PAGE with the indicated antibodies, enlisted below. Molecular weight markers used was Protein Ladder One (Triple-color; Nacalai Tesque, Japan). Anti-HA (3F10) antibody was purchased from Roche. Anti-PROPEP3 antibodies raised in rabbits against both N- and C-terminal fragments of PROPEP3 were described previously (Ross et al., 2014). For detection of extracellular PROPEP3-Venus pool, protein concentrated from the liquid media with Strataclean resin (Agilent Technologies) after filtration was used as an extracellular fraction.

### RNA sequencing and analysis

Five-day-old seedlings grown as described above were pretreated with 0.1 μM Pep1 for 3 days and then exposed to 150 mM NaCl for the indicated times. Three biological replicates were prepared per treatment and genotype. Total RNA was extracted with an RNA extraction kit following the manufacturer’s procedures (NucleoSpin RNA, Machery-Nagel). Each cDNA library was prepared using a TruSeq RNA Library Prep Kit v2 following the manufacturer’s procedures (Illumina, USA). High-throughput sequencing was run by single read 50-bp on a HiSeq2500 platform (Illumina). Raw sequence data were deposited in the DDBJ Sequence Read Archive (accession number DRA004299). Reads were mapped to the TAIR9 Arabidopsis transcriptome database (https://www.arabidopsis.org). The edgeR software package (bioconductor.org.packages/release/bioc/html/edgeR.html) was used for estimation of false discovery rate (FDR) for differential gene expression of raw reads from all 3 biological replicates.

All mRNA variants detected from a gene locus were defined as separate genes in RNA-seq analyses, but assembled and scored for the one gene locus in cross-referencing RNA-seq and ChIP-seq data. For instance, 343 genes were scored as DEGs displaying faster salt induction after Pep1 pretreatment in our RNA-seq analysis, while they were scored as 329 genes corresponding to their loci in the cross-referenced ChIP-seq data. Heatmap was generated with an R-software heat map tool from gplot package (https://cran.r-project.org/web/packages/gplots/) with differentially expressed genes (DEGs) identified using the following cut-off values: FDR <0.05, expression |log2FC ≥1] and Student’s t-test p <0.05. Gene read counts were normalized to RPKM values, and hierarchical clustering was conducted with one minus Pearson correlation complete linkage.

### *In silico* transcription factor binding site analysis

DNA Sequences within 1000 bp upstream of the annotated transcriptional start sites for the indicated genes were examined with the MEME suite 5.0.2 online tool (http://meme-suite.org). Motif enrichment was performed with Local Motif Enrichment Analysis CentriMo with reference to DAP motifs (O’Malley *et al*, 2016) ARABIDOPSIS (Arabidopsis thaliana) DNA database, along the manufacturer’s default settings.

### Bacterial inoculation for salt tolerance assay

*Pseudomonas syringae* DC3000 *ΔhrpS* (Jovanovic *et al*, 2011), *AvrRpm1* (Debener *et al*, 1991), *AvrRps4* (Sohn *et al*, 2009) and *Pseudomonas simiae* WCS417 (Berendsen *et al*, 2015) were grown in NYGB media (5 g/L peptone, 3 g/L yeast extract, 20 mL/L glycerol, pH7.0) supplemented with appropriate antibiotics (rifampicin 25 mg/mL in DMSO, kanamycin 50 mg/mL in deionized distilled water (ddH_2_O), tetracyclin 15 mg/L in ethanol, chloramphenicol 30 mg/mL in ethanol). Overnight bacterial cultures were washed at least twice with 10 mM MgCl_2_ and then adjusted to OD_590_ = 0.002 for spray inoculation. Seedlings were transferred from liquid growth media to agar plates 1 day prior to spray-inoculation. At 6 h after inoculation, seedlings were surface-sterilized twice with 70% ethanol, rinsed twice with autoclaved H_2_O and then transferred to agar media supplemented with or without 175 mM NaCl.

## Acknowledgments

We thank the Saijo Lab members for technical assistance and insightful discussions, and Drs. Yasunari Fujita, Fumiaki Katagiri, Jian-Min Zhou and Cyril Zipfel for published materials. This work was supported in part by the grants from the MEXT of Japan (no. 18H04783 and 18H02467 to Y.S.), grants from SFB670 (Y.S., I.S., J.P.), JST PRESTO (Y.S.), the Asahi Glass Foundation (T.T.) and the Max Planck Society (Y.S., I.S., J.P.), and PhD fellowships from the Japanese Government Scholarship for International Priority Programs (E.L.) and from the Japan Society for the Promotion of Science (T.H.). The RNA sequencing was supported by the Cooperative Research Program of the Genome Research for BioResource, NODAI Genome Research Center, Tokyo University of Agriculture.

## Author contributions

YS conceived the study. EL, KY and YS designed the experiments. EL, YT, KY, TH, HA and TF developed and performed the experiments. EL, YT, KY, HA, KT and TT analyzed the data. IS and JP provided materials and advised on the experiments. EL and YS wrote the manuscript with contributions from the other authors.

